# Early face deprivation leads to long-lasting deficits in cortical face processing

**DOI:** 10.64898/2026.02.04.703627

**Authors:** Saloni Sharma, Margaret S. Livingstone

## Abstract

Experience-dependent plasticity in early sensory cortex is restricted to critical periods that occur early in development. In contrast, when during development higher-order cortical areas show experience-dependent plasticity remains unclear. In the primate ventral visual stream, face-selective cortex emerges with experience during development, providing a powerful system for testing how the timing of experience shapes higher-order representations. Here, we leveraged a unique macaque model of selective face deprivation followed by years of normal face exposure to causally test late plasticity in higher-order visual cortex. We found that face-deprived monkeys acquire face-directed viewing behavior and neural face selectivity after post-deprivation exposure. However, their neural representations retained lasting signatures of early experience, including mixed face and hand selectivity and impaired tuning for face identity, expression, and viewpoint. These findings demonstrate that higher-order visual cortex remains plastic beyond early development, but that the timing of experience fundamentally constrains the refinement of later-acquired neural representations.

## Introduction

How does experience shape the organization of the cerebral cortex, and how does the timing of experience constrain this process? In sensory systems, experience-dependent plasticity is well-established: experiments in cats and monkeys, but also rabbits, hamsters, guinea pigs, songbirds and even humans have shown that altering sensory input during early life can profoundly reshape neural response properties^1–9^. In primates, several lines of recent evidence suggest that the sensitivity to early experience also extends to higher-order cortices including the ventral visual stream, which is organized into regions that respond selectively to specific categories, such as faces^10,11^. First, cortical face selectivity is absent at birth in monkeys and humans, but emerges over the first months of life, coinciding with abundant face exposure^12–14^. Second, extensive early training with non-natural stimuli induces selectivity for those stimuli^15,16^. Third, monkeys that did not see faces during the first year of life fail to develop face domains in inferotemporal cortex (IT), instead showing neural selectivity for hands^12^. Together, these findings indicate that early experience can not only refine maps and create functional domains in early visual areas but also influence the functional properties of higher-level visual neurons.

Importantly, seminal work by Hubel and Wiesel using sensory deprivation demonstrated that such experience-dependent changes in early sensory areas can occur only early in development and are typically long-lasting^1–3^. In contrast, whether higher-order visual cortex remains plastic beyond early development and whether later experience can correct abnormalities induced by early deprivation remains unresolved. Patients who experienced a transient period of early blindness due to congenital cataracts show abnormalities in visually evoked potentials and functional connectivity in higher visual areas^17–19^, yet still show category selectivity in ventral occipito-temporal regions^20^. Similarly, congenitally blind individuals show face-selectivity in temporal cortex to auditory or haptic stimuli^21,22^. These findings have been interpreted as evidence that category selectivity in high-level visual cortex does not require early visual experience^20–22^. However, human studies cannot isolate face-specific deprivation from broader sensory loss, control the timing of experience, or assess the fine-grained neural tuning underlying face processing. Consequently, it remains unclear whether category selectivity acquired after atypical early experience is equivalent to that established during typical development.

To address this question, we studied macaque monkeys reared with selective deprivation of early face experience while preserving all other visual input, followed by prolonged exposure to faces. This paradigm allows a causal test of whether face selectivity can be acquired after early deprivation, and how similar late-acquired selectivity is to that in monkeys with typical early experience. We measured in three previously face-deprived and three typically-reared control monkeys: (1) behavioral gaze biases to faces, hands, objects and face expression using eye-tracking, and (2) category, expression, identity, and viewpoint encoding in neurons in IT cortex with electrophysiological recordings. We found that, behaviorally, the face-deprived monkeys did not differ from controls in their face-looking or face expression preference, though they differed in how they explored faces (upper vs lower bias). Further, IT neurons in face-deprived monkeys showed mixed face and hand selectivity, and persistent impairments in tuning for face identity, expression and viewpoint, in contrast to control monkeys who showed face-only selectivity and robust face expression, viewpoint and identity tuning. These findings demonstrate that neural face selectivity can emerge with later exposure, indicating that higher-order visual cortex remains plastic beyond early development. However, the timing of the experience fundamentally constrains the refinement of late-emerging representations, such that early deprivation leaves long-lasting effects on the functional properties of higher-level visual neurons.

## Results

### Acquisition of face looking behavior

Here we studied three face-deprived monkeys, one of which (B6) had been characterized in prior work from our laboratory^12^, while two additional animals (B14 and B16) were newly included in the present study. As a validation of the deprivation manipulation, we conducted an fMRI localizer in B14 and B16 near the end of the face-deprivation period (see Supplementary Fig. 1 for the experimental timeline). Consistent with prior results from B6, B9, and B10^12^, neither B14 nor B16 showed faces>objects selective regions, instead showing hands>objects selective regions in the parts of IT cortex that would have been face selective in control animals (Supplementary Fig. 2). Likewise, earlier work showed that monkeys B6, B9 & B10 showed a viewing bias for hands more than faces^12^. Building on these established findings, we asked if the face-deprived monkeys recovered from their behavioral hand viewing bias following post-deprivation exposure to faces. To test this, we measured viewing preferences by presenting pairs of faces, hands, and objects while recording gaze position. Fig. 1 shows the results of these experiments conducted at different times after the end of the face-deprivation period, once normal face experience had commenced. 2 months to 1 year after starting exposure to faces, B14 and B16 looked about equally at faces and objects (Fig. 1a, B14: 0.51 ± 0.29, *p* value = 0.89, Wilcoxon signed rank test vs 0.5, B16: 0.54 ± 0.29, *p* value = 0.12) and retained preferential looking at hands (Fig. 1b, B14: 0.46 ± 0.27, *p* value = 0.03, B16: 0.41 ± 0.29, *p* value = 0.00089; Hands (vs objects), B14: 0.53 ± 0.27, *p* value = 0.3, B16: 0.58 ± 0.3, *p* value = 0.0022). But after 2-3 years of exposure to faces, B14 and B16 showed increased face looking (Faces (vs objects), B14: 0.82 ± 0.24, Wilcoxon signed-rank *p* value = 1.1 × 10^-23^, B16: 0.66 ± 0.45, *p* value = 3.6 × 10^-5^; Faces (vs hands), B14: 0.47 ± 0.35, Wilcoxon signed-rank *p* value = 0.4, B16: 0.51 ± 0.47, *p* value = 0.68), though they also still retained some hand bias (Fig. 1c, Hands (vs objects), B14: 0.78 ± 0.27, Wilcoxon signed-rank *p* value = 6.2 × 10^-20^, B16: 0.68 ± 0.44, *p* value = 2.17 × 10^-7^). After 4.5 years of experience seeing faces, B6 looked more at faces (Faces (vs objects) = 0.63 ± 0.34, *p* value = 2.1 × 10^-5^; Faces (vs hands) = 0.83 ± 0.21, *p* value = 7.7 × 10^-21^; Hands (vs objects) = 0.34 ± 0.32, *p* value = 5.8 × 10^-8^) and could not be differentiated from control monkeys (mean Faces (vs objects) = 0.58 ± 0.09, t_4_ = 0.56, *p* = 0.61, Cohen’s d = 0.61; mean Faces (vs hands) = 0.68 ± 0.04, t_4_ = 3.77, *p* = 0.02, Cohen’s d = 4.13; mean Hands (vs objects) = 0.41 ± 0.07, t_4_ = -0.94, *p* = 0.4, Cohen’s d = -1.03; see Supplementary Table 1 for proportion looking time per monkey). Overall, all three face-deprived monkeys showed increasing face viewing preference over time with increased exposure to faces. Next, we investigated how the deprivation and subsequent exposure to faces impacted other aspects of face viewing behavior.

**Fig. 1.**
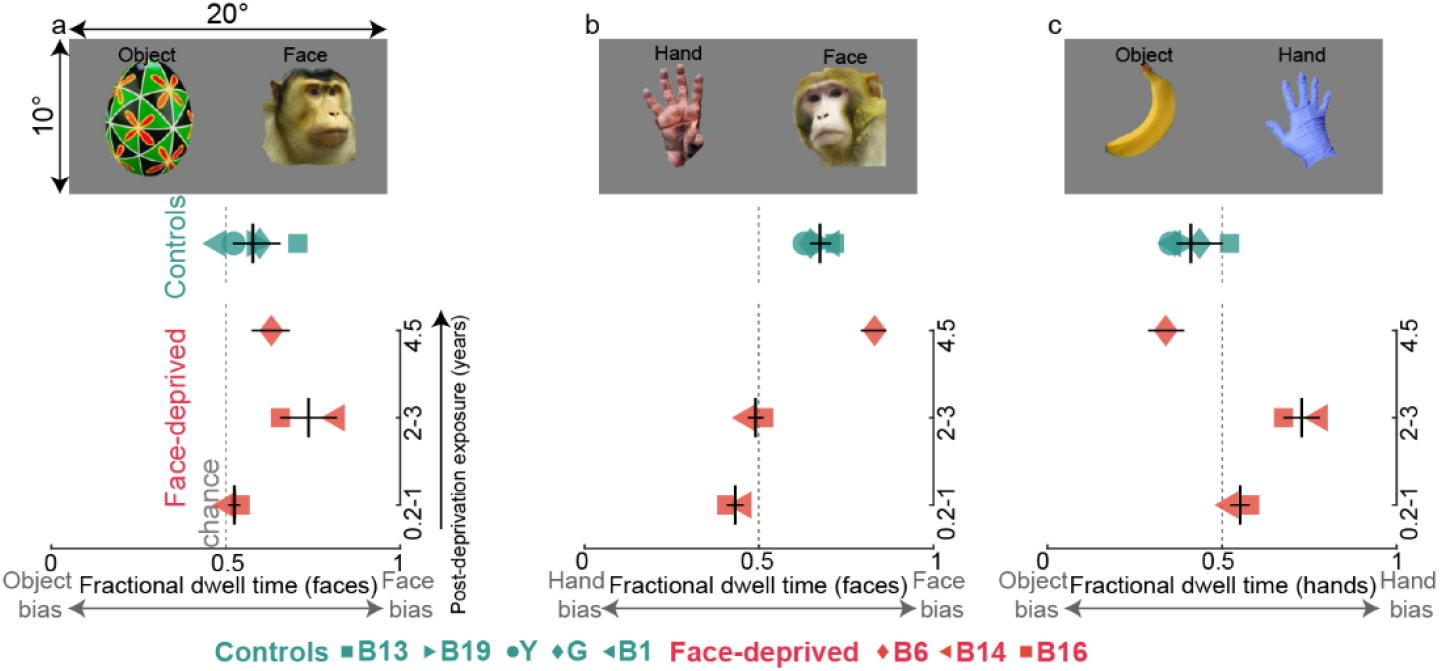
Face-deprived monkeys show some acquisition of face looking behavior after prolonged post-deprivation face exposure. **a**. Viewing behavior shown as fractional dwell time for faces vs objects for control (teal) and face-deprived (red) monkeys. Data for face-deprived monkeys is arranged chronologically by time (on the y-axis) after post-deprivation face exposure (0.2-1yr, 2-3yr, 4.5yr). Individual monkeys are indicated by different symbols. Vertical black line indicates mean and horizontal black line 95% CI across monkeys. For B6 (red diamond), the horizontal black line indicates 95% CI across trials. Grey dotted line indicates chance. **b-c**. Viewing behavior shown as fractional dwell time for faces vs hands (b), and hands vs objects (c) for control (teal) and face-deprived (red) monkeys. Same conventions as in **a**.

### No behavioral expression bias, but different face feature preference

To investigate finer-grained aspects of face processing, we asked how the monkeys looked at different face expressions (Fig. 2a) after a prolonged period of normal face experience. To quantify expression preference, we calculated the proportion of time spent looking at threat vs neutral faces across all trials (Fig. 2a, bottom panel). At the individual level, we found weak effects: the control monkeys looked equally or slightly above chance at threat faces (monkey P: mean = 0.53 ± 0.45, Wilcoxon signed-rank *p* value = 0.14; monkey R = 0.51 ± 0.41, *p* = 0.66; B1 = 0.53 ± 0.45, *p* = 0.37) while the face-deprived monkeys looked equally or slightly more at neutral faces (B6 = 0.52 ± 0.47, *p* = 0.64; B14 = 0.49 ± 0.49, *p* = 0.69; B16 = 0.50 ± 0.39, *p* = 0.87). At the group level, this trend was more evident though still not a strong effect (control mean = 0.52 ± 0.01; face-deprived mean = 0.50 ± 0.01, t_4_ = 2.12, p = 0.10, Cohen’s *d* = 1.73). Next, to assess viewing biases for different parts of a face, we computed the proportion of time the monkeys spent looking at the upper and lower half of each face across all trials (Fig. 2b). Control monkeys looked more at the upper face half (monkey P: mean = 0.56 ± 0.38, Wilcoxon signed-rank *p* value = 0.0027; monkey R = 0.65 ± 0.29, *p* = 1.32 × 10^-14^; B1 = 0.68 ± 0.31, *p* = 2.12 × 10^-28^), while the face-deprived monkeys looked more at the lower half (B6 = 0.03 ± 0.1, *p* = 2.69 × 10^-64^; B14 = 0.29 ± 0.4, *p* = 1.13 × 10^-23^; B16 = 0.19 ± 0.31, *p* = 2.33 × 10^-33^), with a significant difference between the two groups (controls mean = 0.63 ± 0.06; face-deprived mean = 0.17 ± 0.13; t_4_ = 5.47, p = 0.0054, Cohen’s *d* = 4.47). Taken together, these results show that viewing behavior following face deprivation was still abnormal in the face-deprived monkeys in some ways: they largely overlapped with controls in face preference and expression viewing, but systematic differences persisted in within-face feature sampling.

**Fig. 2.**
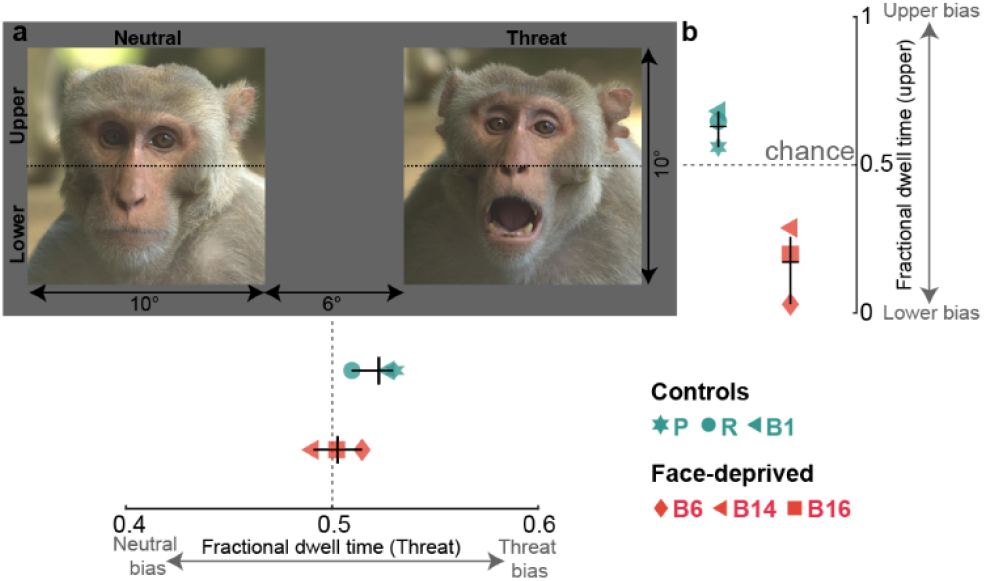
Face-deprived monkeys did not show a looking preference for face expressions, but different face feature preference. **a**. *Top*. Example visual stimuli, of a neutral and a threat face, that were used in the experiment. Images adapted from *Facial Stimuli – Macaques* (Pfefferle, 2020, https://doi.org/10.6084/m9.figshare.13227779.v1), used under CC BY 4.0 license. Each image was shown on both the left and the right to eliminate lateralized viewing biases. *Bottom*. Fractional dwell time for threat and neutral faces for three control (teal) and three face-deprived (red) monkeys. Individual monkeys are indicated by different symbols. Vertical black line indicates mean and horizontal black line 95% CI across monkeys. Grey dotted line indicates chance-level. Note that this experiment was run at a single time point after 4-8 years of normal face exposure, depending on the age of the monkey (see Supplementary Fig. 1 for the experimental timeline). **b**. Fractional dwell time for upper and lower half of the face. Same conventions as in b, except horizontal black line indicates mean and vertical black line indicates 95% CI.

### Acquisition of neural face selectivity, with retained hand selectivity

To test category selectivity of IT neurons, we recorded from electrode arrays implanted chronically in face patches in control monkeys and in face-deprived monkeys in parts of IT that in control monkeys would be face selective^23^. We recorded multiunit spiking activity while the monkeys passively fixated images of faces, hands and objects after 4-8 years of normal face experience. Fig. 3a shows the heatmap of normalized responses to the presented images. Control monkeys showed higher responses to faces compared to hands and objects, while many of the neurons in the face-deprived monkeys showed high responses for both faces and hands compared to objects. We quantified category selectivity for each unit by computing a d’ selectivity index for faces vs objects and a d’ for hands vs objects (Fig. 3b). This index reflects the difference in mean responses between two image categories in standard deviation units; for instance, a face d’ >0 indicates units respond more to faces than to objects, and a hand d’>0 indicates units respond more to hands than to objects. While the control monkeys’ IT neurons are more face selective than hand selective (Monkey P (CIT): *n* = 42, mean face d’ = 1.97, hand d’ = 0.46, t_41_ = 6.23, p = 2.01 × 10^-7^; Monkey P (AIT): *n* = 62, face d’ = 1.17, hand d’ = 0.17, t_61_ = 8.21, p = 1.89 × 10^-11^; Monkey R: *n* = 57, face d’ = 2.4, hand d’ = -0.04, t_56_ = 19.15, p = 2.98 × 10^-26^; B1: *n* = 16, face d’ = 1.99, hand d’ = 0.01, t_15_ = 11.11, p = 1.24 × 10^-8^), the face-deprived monkeys showed mixed face and hand selectivity even after 4-8 years of face experience (B6: *n* = 27, mean face d’ = 0.07, hand d’ = 0.88, t_26_ = -7.36, p =8.13 × 10^-8^; B14: *n* = 41, face d’ = -1.46, hand d’ = 1.65, t_40_ = -1.65, p = 0.11; B16: *n* = 14, face d’ = 1.24, hand d’ = 1.55, t_13_ = -2.77, p = 0.02). Moreover, the face-deprived monkeys showed a positive correlation between face and hand selectivity (*r* = 0.63, p = 2.41 × 10^-10^; Control monkeys: *r* = 0.05, p = 0.51), further indicating mixed face and hand selectivity in face-deprived monkeys (Fig. 3c; see Supplementary Table 2 for correlation values per monkey).

**Fig. 3.**
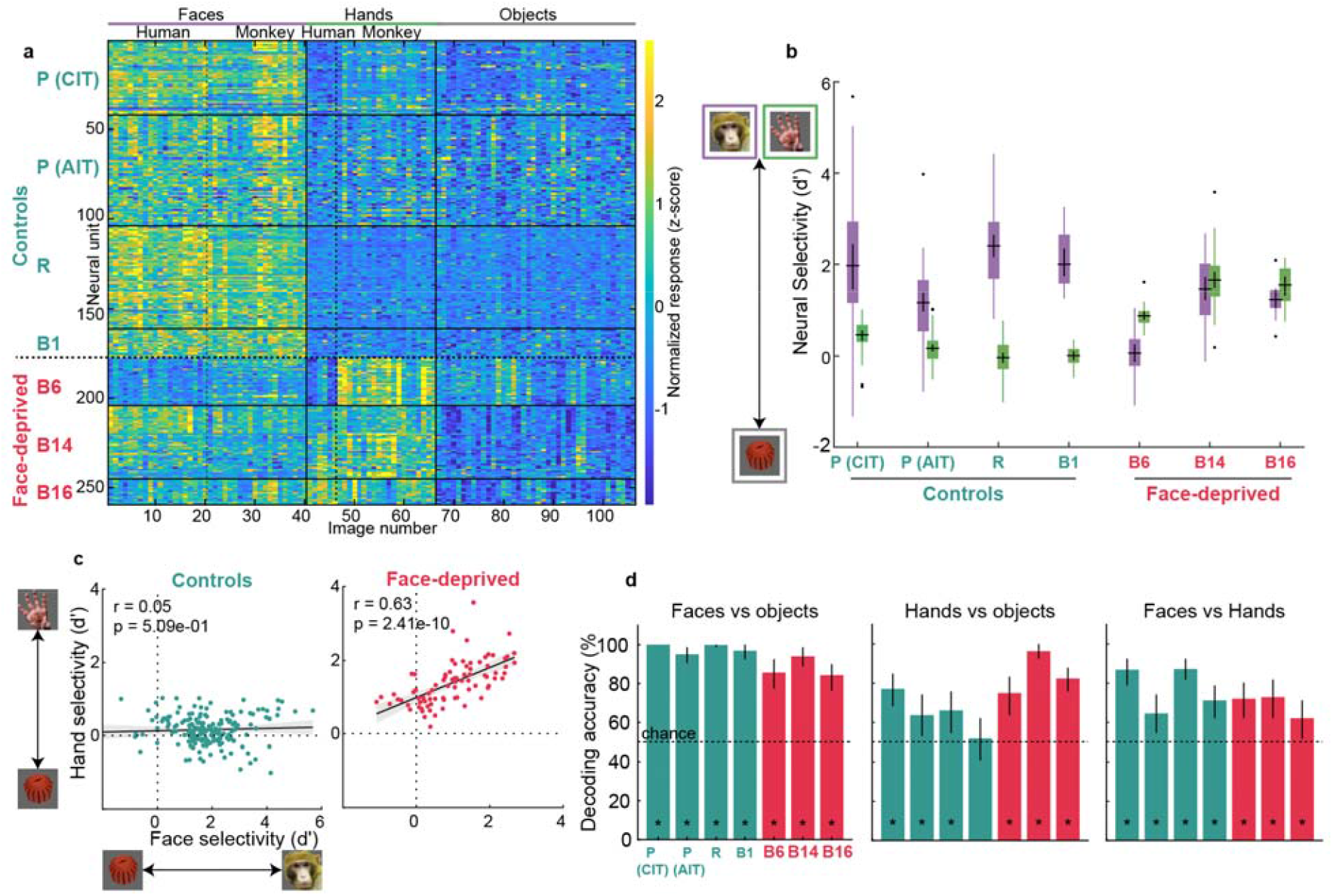
Face deprived monkeys acquire face selectivity, while retaining abnormally strong hand selectivity. **a**. Response heatmap shows responses of individual neural units (rows) to 40 faces, 26 hands, and 40 inanimate non-face objects (columns) for 3 control and 3 face-deprived monkeys. Responses were normalized (z-scored) per site across images. Color scale limits were set to the 1^st^ and 99^th^ percentiles of all normalized responses to reduce the influence of extreme values. **b**. Neural selectivity (d’) for faces vs objects (purple boxplots) and hands vs objects (green boxplots) for 3 control and 3 face-deprived monkeys. The black central horizontal line shows the mean response, the black central vertical line indicates confidence intervals, and the bottom and top edges of the box indicate the 25th and 75th percentiles, respectively. Whiskers (vertical lines in color) show the most extreme data points not considered outliers. Outliers are indicated by black crosses. **c**. Scatterplot showing the correlation between face selectivity (face d’) on the x-axis and hand selectivity (hand d’) on the y-axis. Each dot indicates a neural unit in control monkeys (left panel; teal dots; n = 177, pooled across 3 monkeys) and face-deprived monkeys (right panel; red dots; n = 82, pooled across 3 monkeys). The black line indicates an ordinary least squares linear regression fit, with shaded 95% confidence intervals error bands. The values on the top left corner indicate Pearson’s correlation (*r*) and the corresponding *p*-value. **d**. Neural population-level decoding accuracy for category pairs (faces vs objects, hands vs objects, faces vs hands) in 3 control (teal) and 3 face-deprived (red) monkeys per monkey. Error bars indicate 95% bootstrap confidence intervals across 10,000 iterations (see Methods). Dotted line indicates chance level (50%). Asterisks indicate monkeys with decoding significantly above chance (lower CI bound > chance).

A decoding analysis aimed at calculating how much category information can be extracted from neural population responses showed that neuronal populations from both control and face-deprived monkeys perform significantly above chance at decoding face vs object or hand vs object categories (Fig. 3d; chance level = 50%). However, face vs object decoding accuracy was higher in control monkeys (mean = 98.1 ± 1.6%) than in face-deprived monkeys (mean = 88 ± 5.27%; t_4_ = 3.17, p = 0.034, Cohen’s d = 2.59). Face vs hand decoding accuracy was higher in control monkeys (mean = 78.02 ± 8.23%) than in face-deprived monkeys (mean = 68.99 ± 5.99%; t_4_ = 1.54, p = 0.2, Cohen’s d = 1.25). Interestingly, IT neurons in face-deprived monkeys decoded hands vs objects better (mean = 84.5 ± 10.8%) than control monkey IT neurons (62.8 ± 9.7%), indicating that control monkeys represent hands more similarly to objects, whereas face-deprived monkeys represented hands as more distinct from objects (t_4_ = -2.6, p = 0.061, Cohen’s d = -2.12). Supplementary Fig. 4 shows multidimensional scaling (MDS) of neural population responses to faces, hands, and objects in face-deprived and control monkeys. In control monkeys faces cluster separately from an overlapping cluster of hands and objects. In the face-deprived monkeys, faces and hands clustered together, separate from objects. Overall, these results indicate that the face-deprived monkeys developed mixed selectivity for hands and faces after post-deprivation exposure to faces. That is, the IT neurons retained the hand selectivity they developed during face deprivation but also acquired face selectivity. This contrasts with control monkeys, who showed a strong and stable face selectivity (see Supplementary Fig. 3 to compare neural selectivity over time). This divergence from control monkeys raises an important question: how does early face deprivation impact core IT neuron properties^24–26^, such as face expression and identity tuning?

### Impaired face expression and identity tuning

We investigated IT neural selectivity for face expressions, by computing a threat d’, i.e., a d’ higher than 0 indicates higher responses for threat vs neutral faces and lower than 0 indicates higher responses for neutral vs threat faces (Fig. 4b). We found that control monkey neurons showed higher responses for threat compared to neutral faces (monkey P: *n* = 51, threat d’ = 0.21 ± 0.35; R: *n* = 62, d’ = 0.46 ± 0.33, B1: *n* = 10, d’ = 0.25 ± 0.59), while neurons in face-deprived monkeys did not show such selectivity (monkey B6: *n* = 31, threat d’ = 0.003 ± 0.28; B14: *n* = 55, d’ = -0.17 ± 0.3, B16: *n* = 18, d’ = -0.10 ± 0.38). We further found that face selectivity was positively correlated with threat selectivity in control monkeys (Fig. 4c; Pearson’s *r =* 0.19, *p* = 0.037), whereas face and threat selectivity were negatively correlated in face-deprived monkeys (Pearson’s *r =* -0.42, *p* = 8.56 × 10^-6^; see Supplementary Table 3 for correlation values per monkey). We asked if we could decode face expression information from IT units (Fig. 4d) and found that control-monkey neurons demonstrated a higher decoding accuracy (mean = 69.4 ± 2.14%; chance level = 50%) than neurons in face-deprived monkeys (mean = 54.1 ± 4.15%; t_4_ = 5.7, p = 0.0047, Cohen’s d = 4.65). We found a similar trend for identity decoding, with control-monkey neurons performing better than neurons in face-deprived monkeys (Fig. 4e; control-monkeys mean = 30.1 ± 17.71%; face-deprived = 13.77 ± 6.94%; chancel level = 5.55 %; t_4_ = 1.49, p = 0.21, Cohen’s d = 1.21). Overall, these results indicate impaired face expression and face identity representation in IT neurons from face-deprived monkeys. Next, we investigated if another hallmark property of IT face cells, viewpoint tuning^27,28^, is also disrupted in neurons in the face-deprived monkeys.

**Fig. 4.**
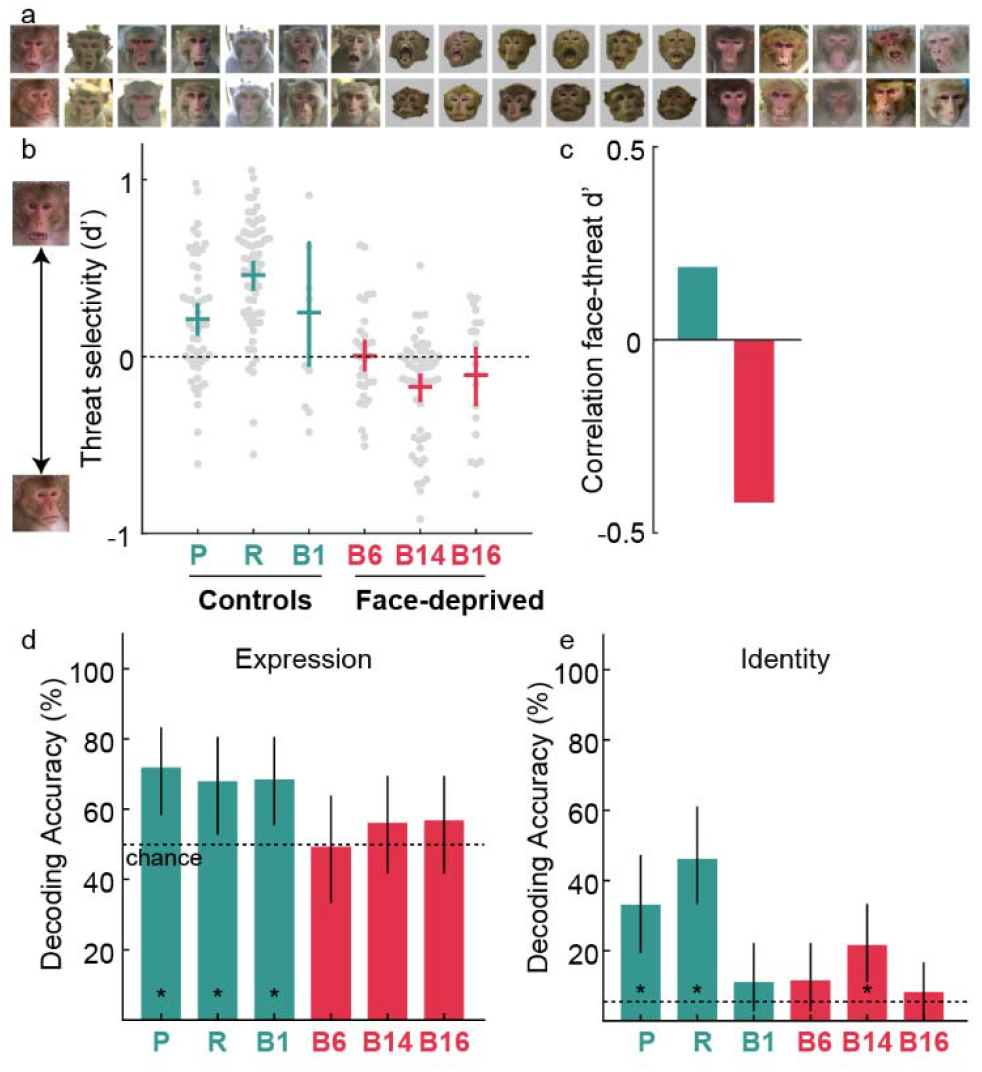
IT neurons in face-deprived monkeys showed impaired face expression and identity tuning. **a**. The visual stimuli, depicting threat (top) and neutral (bottom) faces, that were used in the experiment. Images adapted from *Facial Stimuli – Macaques* (Pfefferle, 2020, https://doi.org/10.6084/m9.figshare.13227779.v1), used under CC BY 4.0 license. **b**. Average threat selectivity (d’) for 3 control (teal) and 3 face-deprived (red) monkeys. The central horizontal line shows the mean selectivity, and the central vertical line indicates confidence intervals. Beeswarm plots show the threat selectivity of each individual unit as a gray dot. **c**. Correlation (Pearson’s *r*) between face and threat selectivity for each group, computed by pooling units across the 3 control (teal) and 3 face-deprived (red) monkeys. The correlation per monkey and the corresponding p value is shown in Supplementary Table 3. **d**. Neural population-level decoding accuracy for the 2 face expressions (threat and neutral), shown per monkey. Error bars indicate 95% bootstrap confidence intervals across 10,000 iterations. Dotted line indicates chance level (50%). Asterisks indicate monkeys with decoding significantly above chance (lower CI bound > chance). **e**. Neural population-level decoding accuracy for 18 face identities (shown in a) per monkey. Same conventions as in d, except chance level = 5.55%.

### Reduced viewpoint and identity tuning

To ask if we could decode viewpoint/identity information from IT neurons, we presented the monkeys with 7 different identities and 5 different viewpoints (Fig. 5a). Neurons in control monkeys performed better than neurons from face-deprived monkeys for viewpoint decoding, though both groups were above chance (control monkey neurons mean = 63.19 ± 21.53%; face-deprived = 39.88 ± 3.48%; chancel level = 20%; t_4_ = 1.85, p = 0.14, Cohen’s d = 1.51). We observed a similar trend for identity decoding, with neurons from control monkeys performing better than neurons from face-deprived monkeys, though both were above chance (control neurons monkeys mean = 71.33 ± 29.35%; face-deprived = 32.88 ± 2.34%; chancel level = 14.29%; t_4_ = 2.26, p = 0.087, Cohen’s d = 1.85). When identity decoding was reduced to three cardinal viewpoints (full left, front, full right), decoding accuracy in face-deprived IT slightly decreased (31.51 ± 1.14%). Importantly, the lower bound of the confidence intervals dropped to chance level (14.3%), indicating less robust identity representation across views. In contrast, control monkey neurons maintained high accuracy and above-chance confidence intervals under both conditions. Overall, these results indicate that even after 4-8 years of exposure to faces, the face-deprived monkeys did not fully gain neural properties typically associated with IT face cells.

**Fig. 5.**
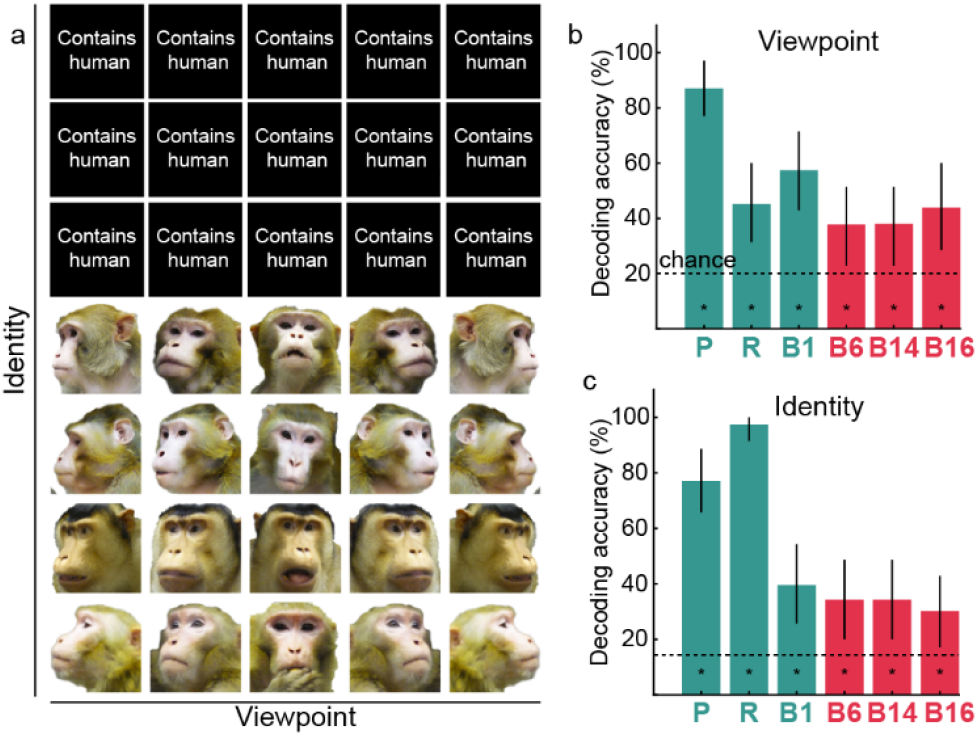
Face-deprived monkeys showed reduced viewpoint and identity tuning. **a**. The visual stimuli, depicting 7 identities and 5 viewpoints, that were used in the experiment. Images were taken in the lab. **b**. Neural population-level decoding accuracy for the 5 viewpoints for 3 control (teal) and 3 face-deprived (red) monkeys. Error bars indicate 95% bootstrap confidence intervals across 10,000 iterations. Dotted line indicates chance level (20%). Asterisks indicate monkeys with decoding significantly above chance (lower CI bound > chance). **c**. Neural population decoding accuracy for the 7 face identities per monkey. Same conventions as in b, except chance level = 14.29%.

## Discussion

In this study, we investigated long-lasting effects of early face deprivation on face processing and the scope of acquisition of face processing after the deprivation period. We conducted behavioral and neurophysiology experiments with three face-deprived monkeys after prolonged face exposure and three typically raised control monkeys. We found that, behaviorally, face-deprived monkeys largely overlapped with controls in face preference and expression viewing, though the face exploration patterns (upper vs lower) differed from controls. Neurally, we found that IT neurons in face-deprived monkeys showed (1) mixed face and hand selectivity; (2) impaired face expression and identity tuning and (3) reduced viewpoint and identity tuning, in contrast to control monkeys who showed face-only selectivity and robust face expression, viewpoint and identity tuning. This indicates that while coarse category selectivity can emerge after deprivation, early experience left a lasting imprint on cortical organization. Overall, our findings show that early visual experience can fundamentally shape neural representations across the visual system, including higher-order regions such as inferotemporal cortex. Further, although later exposure can alter behavioral consequences during naturalistic viewing, it did not fully produce typical neural organization.

Our finding of late acquisition of neural face selectivity helps reconcile seemingly disparate results from human studies of abnormal early visual experience. Patients treated for congenital cataracts have shown face-selective responses in the ventral visual stream despite their early visual deprivation^20^. Furthermore, congenitally blind individuals show category-selectivity in temporal cortex to auditory or haptic stimuli^21,22^. These studies have been interpreted as evidence that category selectivity in high-level visual cortex does not require visual experience^20–22^. Our results are consistent with the observation that face selectivity can be acquired later in life, even after an extended period of atypical early visual experience but provide a more direct test of its dependence on experience. Our paradigm selectively eliminated face experience while preserving all other visual experience and allowed before-and-after measurements in the same animals. This early face-specific deprivation prevented the development of face-category selectivity at the time normally reared monkeys acquire it, yet face selectivity reliably emerged following later exposure to faces. Thus, our results demonstrate that face selectivity in inferotemporal cortex depends on the availability of face experience. Indeed, neither congenitally blind nor cataract-reversal populations lack face experience altogether: blind individuals have extensive tactile and auditory experience with faces, and cataract-reversal patients in ref^20^ experienced only transient neonatal deprivation and were tested years after extensive face exposure. Preserved face selectivity in these groups therefore does not rule out an experience-dependent account of the emergence of category selectivity from later experience, as we observe here in our results.

Our findings raise an important question: why does face selectivity emerge in the same stereotypical location in the face-deprived monkeys after post-deprivation face exposure? We previously observed strong hand selectivity in face-deprived monkeys during the deprivation period, in locations that would be face selective in normally reared monkeys^23^. We suggested that this abnormal selectivity arose because, in the absence of faces, these young monkeys must look extensively at their own hands manipulating objects and food and climbing around their environment, as well as at the often present and socially important hands of care staff. Thus, these regions correspond to portions of IT that represents the central visual field, and which are preferentially driven by foveated input and thus particularly sensitive to the statistics of visual experience^15,29–33^. Our current study provides further evidence for this account: following the introduction of faces, foveation in the face-deprived monkeys shifts toward faces, allowing face-selective responses to emerge in the same part of IT that was previously shaped by hand viewing. This account is consistent with human developmental findings showing that, as gaze and visual priorities shift with development, ventral temporal regions are repurposed to represent faces, limbs and text^34–38^. Further support comes from findings beyond faces, showing that intensive early experience with specific stimulus classes can give rise to category selectivity in ventral temporal cortex, even for ethologically arbitrary stimuli. For example, humans show selectivity for Pokemon characters following intensive childhood experience, and monkeys develop selectivity for extensively trained symbols^15,32,39^.

Our finding of the later emergence of face selectivity suggests that high-level visual areas remain malleable well beyond early infancy, extending at least into later pre-pubertal years^40^. Work by Hubel and Wiesel provides a useful framework for interpreting these findings: closing one eye (monocular deprivation) early in development shifted neuronal selectivity to the open eye, but this effect diminished with age and became undetectable after 1 year^41^. If one eye was closed for a short period, then that eye was opened and the other eye closed, inputs from the later opened eye could overwrite the earlier opened eye, but within restricted developmental windows: up to 3 weeks for the magnocellular inputs to layer 4Cα, and 6 weeks for parvocellular inputs to layer 4Cβ. These results indicate that different parts of the brain have different critical periods during which experience can modify selectivity^8,20,42,43^. In this context, the ability of monkeys to acquire face selectivity after a year of deprivation indicates that the critical period for plasticity in higher visual areas must extend later than the critical period for V1.

Importantly, while this extended plasticity reflects a prolonged critical period, it should be distinguished from selectivity acquired after this window, which is more constrained. That is, despite the acquisition of face selectivity post-deprivation, IT neurons in the face-deprived monkeys demonstrated mixed hand and face selectivity rather than a fully categorical organization, and atypical finer-grained aspects of face representations. This persistence of earlier selectivity contrasts with typical development, in which later-acquired categories, if acquired within the critical period, tend to dominate cortical territory^38,44^. One possibility is that later face experience occurs near the end of the IT critical period, and plasticity is diminished. Alternatively, early-established selectivity may simply be permanent: even with the earliest, longest monocular deprivations, Hubel and Wiesel found single-neuron responsiveness to the deprived eye^41^. Thus, incomplete acquisition must reflect the lasting consequences of early-established selectivity that continues to constrain subsequent refinement by later experience. Consistent with this view, studies of literacy show that learning to read in adulthood leads to text selectivity emerging within ventral temporal cortex that retains greater responsiveness to previously established categories, whereas earlier acquisition during childhood results in more fully specialized representations^39,44^. In this way, we suggest that text learning in adult humans and late face exposure in monkeys illustrate how later experience only partially overwrites the effects of earlier experience.

It is notable that the face-deprived monkeys showed largely similar face-looking behavior to control monkeys after prolonged face exposure, despite having had no experience with faces during the first year of life. One possibility is that free-viewing behavior reflects a coarser threshold for detecting abnormalities in face processing. Behavioral differences may therefore emerge more clearly under tasks that require explicit judgments about face identity or expression. Importantly, free-viewing paradigms provide a powerful and ecologically valid assay of visual behavior and can reveal how abnormalities in neural circuitry relate to behaviorally meaningful measures such as gaze patterns^12,45–47^. Indeed, our findings complement prior task-based evidence showing that aspects of fine-grained face processing were comparable to control monkeys within a year after deprivation ends^48^. At the same time, our findings extend those results by revealing persistent differences in how face-deprived and control monkeys sample faces. Thus, behavior becomes typical at the level of coarse preferences but remains atypical in finer-grained sampling strategies.

In conclusion, in this study, we leveraged a unique face-deprivation and post-deprivation exposure paradigm to study the role of visual experience and developmental timing in shaping face-selective cortex. We found that neuronal face-category selectivity and face-directed behavior can be acquired following prolonged face exposure, even when that exposure occurs only later in life. However, this late acquisition was not identical to the face processing of animals with normal early face experience: neural representations retained signatures of early deprivation, including mixed face and hand selectivity and impaired tuning for fine-grained face properties of expression and identity. Taken together, our findings demonstrate that higher-level visual cortex plasticity extends well beyond early infancy, suggesting that different regions in the brain may have different critical periods. At the same time, later plasticity was not unconstrained: early-established selectivity was not simply overwritten by later experience but continued to shape (and in some cases limit) the organization and refinement of subsequent representations. Overall, our findings help situate cortical plasticity along a developmental continuum, showing how experience-dependent reorganization differs between early and later life, a distinction reminiscent of language acquisition^49^: a new language can be learned in adulthood but rarely reaches the fluency and precision achieved when learned early in development.

## Methods

All procedures were approved by the Harvard Medical School Institutional Animal Care and Use Committee (protocol #ISO00001049) and conformed to NIH guidelines provided in the Guide for the Care and Use of Laboratory Animals.

### Subjects and array location

A total of ten monkeys (eight male Macaca mulatta, 5-13 kg and two male Macaca nemestrina, 13-15kg) participated in the experiments. Three of the monkeys were face-deprived, i.e., they were prevented from seeing faces for the first years of their life (1, 1.2 and 1.8 years for B6, B14 and B16, respectively). This was achieved by hand-rearing the monkeys by lab staff members wearing welder’ masks (similar procedure as described in Arcaro et al, 2017^12^; B6 in that paper is the same monkey as B6 in the current paper). In this period, the only face exposure that B14 had was in the fMRI scanning sessions conducted at 130, 143, 193, and 214 days old and B16 at 382 days old. The results shown in Supplementary Fig. 2 correspond to scanning conducted at 214 and 382 days old for B14 and B16 respectively. Both monkeys also saw a few face images during electrophysiology sessions that were run daily for less than an hour a day (of which face viewing time per day was ∼2–7 s) for three months before the end of the deprivation period. Six of the ten monkeys participated in the electrophysiology experiments and were chronically implanted with floating microelectrode arrays (FMA; 32 channels, MicroProbes, Gaithersburg, MD or 128 channels, NeuroNexus, Ann Arbor, MI), microwire bundles (64 channels; MicroProbes) or Neuropixel arrays (45mm; IMEC, Leuven, Belgium). The target locations for the array placement in central or anterior IT was determined based on fMRI localization or anatomical markers (“bumps”^23^) in the lower bank of the superior temporal sulcus. For the face-deprived monkeys, B14 and B16 had arrays in central IT, and B6 in anterior IT. For the three control monkeys, monkey P had an array in central IT and Monkey R and B1 in anterior IT. For one of the experiments (Fig. 1), monkey P had arrays in both central and anterior IT.

### Visual experiments

For the fMRI experiments, the monkeys sat prone in a plastic chair, and visual stimuli were projected 52 cm from the monkey’s eyes, onto a screen at the end of the scanner bore. All the images were 20° in size. The monkeys were rewarded with juice for looking at the center of the screen. Gaze direction was monitored using an infrared eye tracker (ISCAN, Woburn, MA, http://www.iscaninc.com/). For the electrophysiology and behavioral experiments, the monkeys sat upright in a plastic monkey chair and faced an LCD display screen 53 cm in front of the monkey. Monocular eye-tracking signals were acquired at 1lllkHz from an ISCAN infrared eye tracker without digital smoothing or filtering. Analog outputs from ISCAN trackers were sampled at a higher rate (1lllkHz) than the camera frame rate (120lllHz). All experiments were run using the MonkeyLogic software (https://monkeylogic.nimh.nih.gov/), which also monitored and recorded eye-tracking signals. For the electrophysiology experiments, the images (6-8°) were presented in on a grey background at the rate of 100-183ms on, 183-200ms off at the center of the array’s mapped receptive field. The monkeys fixated a red dot (0.2 x 0.2°) in the center of the screen and received continuous juice reward for maintaining fixation. For the behavioral experiments, monkeys performed a free-viewing task where images were presented at a size of 20 x 10° for the category experiment and 26 x 10° for the face expression experiment. The monkeys initiated a trial by fixating a red central dot (0.2°), after which image pairs were presented for 1.5, 2 or 3 seconds in different sessions. The images were presented in a pseudorandom order and repeated when all images had been shown once. Monkeys were rewarded at fixed intervals with a drop of juice for maintaining their gaze within a window around the image.

### Stimuli

For the fMRI experiment in B16, 20 monkey faces, monkey hands and objects centered on a pink-noise background were used (see Supplementary Fig.1 of ref^12^ for examples). For B13 and B14, an additional block of human faces was also included. For the electrophysiology experiments, to quantify neural category selectivity, the experiments included 40 faces (20 human faces, 20 monkey faces), 40 non-faces (20 familiar, 20 nonfamiliar) and 26 hands (6 human, 20 monkey) on a white background (also used in refs. ^50,51^). The 20 non-familiar images of nonfaces are from ref. ^52^, while the 20 images of nonfaces familiar to the monkeys, human and monkey face images were taken in our lab. For the behavioral experiment that measured category preference, we selected a subset of 24 images from this same stimulus pool: 8 faces (4 monkeys and 4 humans), 8 hands (4 monkeys and 4 humans) and 8 inanimate objects (4 familiar and 4 unfamiliar). Each image appeared on the left of the screen with every other stimulus on the right, and vice versa, ensuring that each unique pair was shown once in each left/right configuration, for a total of 552 image pairs. To probe neural threat selectivity and face expression tuning, we used images obtained from publicly available Facial Stimuli – Macaques dataset^53^, which included 18 monkey identities and 2 face expressions (Fig. 4a). A subset of these images was used in the behavioral experiment probing face expression preference, in which eight identities (each with a neutral and a threat expression) were selected. Each neutral expression was paired with every threat expression, yielding 64 unique cross-expression pairs. Each pair was presented in both left-right configurations (neutral on the left/threat on the right and vice versa), resulting in a total of 128 image pairs. Finally, to probe viewpoint and identity tuning (Fig. 5a), we used images depicting 7 different face identities (3 human, 4 monkey) and 5 viewpoints, which were taken in our lab.

### Scanning

Imaging was performed on a 3 T Tim Trio scanner with an AC88 gradient insert using custom four-channel surface coils (A. Maryam, Martinos Imaging Center). Acquisition parameters were TR = 2 s, TE = 13 ms, flip angle = 72°, iPAT = 2, 1-mm isotropic voxels, matrix 96 × 96, and 67 contiguous sagittal slices. To increase SNR^54^, a contrast agent (monocrystalline iron oxide nanoparticles (MION; Feraheme, AMAG Pharmaceuticals, Cambridge, MA) was injected into the saphenous vein just before scanning (12 mg/kg). Because MION inverts the functional signal, we plot negative signal changes as positive for readability. During scanning, monkeys were alert with the head stabilized non-invasively using a foam-padded helmet with chin strap and a bite bar that also delivered juice rewards. Animals sat in a primate chair modified for small monkeys and positioned either semi-upright or in a sphinx posture; bodies and limbs could move freely while the padded helmet maintained a forward-facing head position. Blocks were organized by image category and presented in counterbalanced order. Each image appeared for 0.5s within 20s blocks, separated by 20s of a neutral gray screen.

## Data analysis

### *fMRI* data

Preprocessing and analysis were conducted using statistical parametric mapping (SPM12; https://www.fil.ion.ucl.ac.uk/spm/) and JIP Toolkit (https://www.nmr.mgh.harvard.edu/∼jbm/jip/). Functional series were realigned and nonrigidly coregistered in JIP to each monkey’s high-resolution anatomical template, resampled to 1-mm isotropic voxels, and spatially smoothed with a 2-mm FWHM Gaussian kernel. Condition-specific MION responses were estimated voxelwise with a general linear model (GLM) using previously described procedures^54,55^. Briefly, a MION hemodynamic response function was convolved with boxcar regressors for each stimulus condition, and six motion parameters (three translations and three rotations) were included as nuisance regressors. For visualization, whole-brain SPM T-maps were rendered on an inflated NMT standard template (v1.2^56^) using AFNI (SUMA; https://afni.nimh.nih.gov/Suma). We used 22 runs from B13, 30 runs from B14, and 26 runs from B16 for the analysis.

### Behavioral data

Eye position was sampled at 1 kHz during image presentation. For each millisecond, gaze was labeled “on-image” if the eye position (*x, y*) fell within the image bounds and outside a central vertical gap. For a given two-way contrast (A vs B, e.g., faces vs objects, faces vs hands, hands vs objects, threat vs neutral, or upper vs lower), on-image samples in each trial were classified into *T*_A_ or *T*_B_ using predefined criteria (e.g., left vs right: x>0 vs x<0; upper vs lower: y>0 vs y<0). We defined the *fractional dwell time to A* as

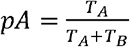

For each monkey, we summarized *pA* as mean ± standard deviation across trials. Values >0.5 indicate a bias toward A; 0.5 indicates no bias; <0.5 indicates a bias toward B (samples off-image, in the gap, or outside A/B were excluded from the denominator). Deviation from chance (0.5) was tested with a two-sided one-sample Wilcoxon signed-rank test on p−0.5 across trials.

### Neural data

#### Firing rates

Neural signals were amplified and sampled at 40 kHz using a data acquisition system (OmniPlex, Plexon, Dallas, TX) for the floating microelectrode and the microwire brush arrays. Multi-unit spiking activity was detected using a threshold-crossing criterion. The Neuropixels data were acquired at 30 kHz using SpikeGLX software (Janelia Research) and preprocessed with CatGT for global-average referencing. Multi-unit activity envelopes (MUAe) were computed using custom code following established methods. The resulting MUAe provides an instantaneous measure of aggregate spiking activity close to the electrode, while being independent of arbitrary spike-detection thresholds. The neural response was defined as the spike rate in a 150ms time window starting at 50-100ms after the image onset, depending on the monkey. The response onset was determined by visually inspecting the average time course of neural activity averaged across all units per monkey. The firing rates per neural site were trial averaged per image.

#### Response reliability

The selection criterion for inclusion of neural units in all analyses was determined by the firing-rate reliability per neural site. To assess unit reliability, trials for each image were randomly split into two halves. For each set, the neural responses were averages across trials to create two response vectors, and the correlation between the two response vectors was computed. This procedure was repeated 100 times with different random splits, and the resulting correlation values were averaged to obtain a single average correlation r. The reliability ρ was computed by applying the Spearman-Brown correction to the average correlation r:

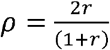

Only neural units with a split-half reliability ρ >0.4 were included in further analysis. The number of units that were included per monkey in each experiment are provided in the main text.

#### Selectivity index

We quantified face selectivity using a d’ index, which compared trial-averaged responses to faces and to non-faces:

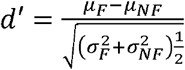

where μF and μNF are the across-stimulus averages of the trial-averaged responses to faces and non-faces, and σF and σNF are the across-stimulus standard deviations. We used the same d’ metric to quantify hand and threat selectivity, which was computed based on neural responses to hands vs objects, or threat vs neutral images, respectively.

#### Decoding analysis

To assess whether information about category, expression, viewpoint and identity could be extracted from neural activity, we used a split-half decoding scheme with bootstrapped accuracy estimation^57^. Neural responses for each stimulus were arranged into a matrix of neurons x trials x stimuli. To control for unequal trial counts, we randomly subsampled trials to match the minimum number of trials available for any stimulus, ensuring equal representation while retaining trial-to-trial variability. For each monkey, we performed 10,000 bootstrap iterations. In each iteration, trials for each stimulus were randomly split into two halves. Reponses within each half were averaged to produce one training and one testing vector per stimulus. Training vectors were z-scored across stimuli, and the same mean and standard deviation were applied to normalize the corresponding test vectors. Linear support vector machine (SVM) classifiers with error-correcting output codes (ECOC) were trained to decode either category/face expression (binary classification of face vs objects, face vs hands, hands vs objects or threat vs neutral) or identity/viewpoint (multiclass classification of the unique identities or the five viewpoints). Decoding accuracy was computed as the proportion of correctly classified stimuli in the test set for each bootstrap iteration. For each monkey, mean accuracy and 95% confidence intervals (2.5^th^ – 97.5^th^ percentiles across bootstrap iterations) were calculated. Decoding performance was considered significantly above chance if the lower bound of the confidence interval exceeded the relevant chance level (computed by calculating 1/n^*^100; leading to chance = 50% for expressions and categories;1/n identities or 20% for 5 viewpoints for identity/viewpoint decoding).

#### Multidimensional scaling (MDS) analysis

To visualize the representational geometry of neural population responses in inferotemporal cortex, we performed metric multidimensional scaling (MDS) separately for each monkey. Neural responses were defined as trial-averaged firing rates per neural site for each image. Images belonging to the face, hand, and object categories were selected based on stimulus annotations. Each image was represented as a population response vector across all recorded neurons, and pairwise dissimilarities between images were computed using Euclidean distance without normalization across neurons. The resulting image to image distance matrix was embedded into two dimensions using metric MDS (MATLAB mdscale), and the resulting coordinates were plotted with points colored according to stimulus category.

#### Statistical inference

P-values were calculated using paired t-tests when comparing differences between conditions within monkeys, and unpaired t-tests when comparing between-group face deprived vs control monkey differences. To account for the small sample size (n = 3 per group), we also report effect sizes (Cohen’s d)^58^.

## Supplementary Figures and Tables

**Supplementary Table 1.** Behavioral preference for object category per monkey. Table showing fractional dwell time (FDT) and the corresponding p-value (Wilcoxon signed-rank test vs 0.5) per monkey (related to Fig. 1).

| Control Monkey | Face vs objects |  | Faces vs Hands |  | Hands vs Objects |  |
| --- | --- | --- | --- | --- | --- | --- |
|  | FDT | p-value | FDT | p-value | FDT | p-value |
| B1 | $0.48 \pm 0.36$ | 0.79 | $0.71 \pm 0.3$ | 0.022 | $0.36 \pm 0.35$ | 0.1 |
| B13 | $0.71 \pm 0.41$ | $2.32 \times 10^{-7}$ | $0.72 \pm 0.41$ | $8.65 \times 10^{-8}$ | $0.52 \pm 0.46$ | 0.59 |
| G | $0.59 \pm 0.29$ | 0.00018 | $0.65 \pm 0.26$ | $1.96 \times 10^{-9}$ | $0.44 \pm 0.29$ | 0.012 |
| B19 | $0.58 \pm 0.29$ | 0.00015 | $0.67 \pm 0.29$ | $3.27 \times 10^{-12}$ | $0.38 \pm 0.33$ | $6.09 \times 10^{-6}$ |
| Y | $0.52 \pm 0.36$ | 0.57 | $0.63 \pm 0.35$ | 0.00019 | $0.35 \pm 0.35$ | $3.27 \times 10^{-5}$ |

**Supplementary Table 2.** Correlation between face and hand selectivity per monkey. Table showing Pearson’s correlation *r* between face and hand d’ for each monkey (related to Fig. 2c).

| Monkey | r | p |
| --- | --- | --- |
| Controls |  |  |
| P (CIT) | 0.42 | 0.0058 |
| P (AIT) | -0.22 | 0.084 |
| R | 0.14 | 0.32 |
| B1 | -0.32 | 0.22 |
| Face-deprived |  |  |
| B6 | 0.01 | 0.95 |
| B14 | 0.43 | 0.0046 |
| B16 | 0.43 | 0.13 |

**Supplementary Table 3.** Correlation between face and threat selectivity per monkey. Table showing Pearson’s correlation *r* between face and threat d’ for each monkey (related to Fig. 4b).

| Monkey | r | p |
| --- | --- | --- |
| Controls |  |  |
| P | 0.32 | 0.02 |
| R | 0.11 | 0.39 |
| B1 | -0.32 | 0.37 |
| Face-deprived |  |  |
| B6 | -0.29 | 0.12 |
| B14 | -0.58 | $4.09 \times 10^{-6}$ |
| B16 | -0.097 | 0.7 |

**Supplementary Figure 1.**
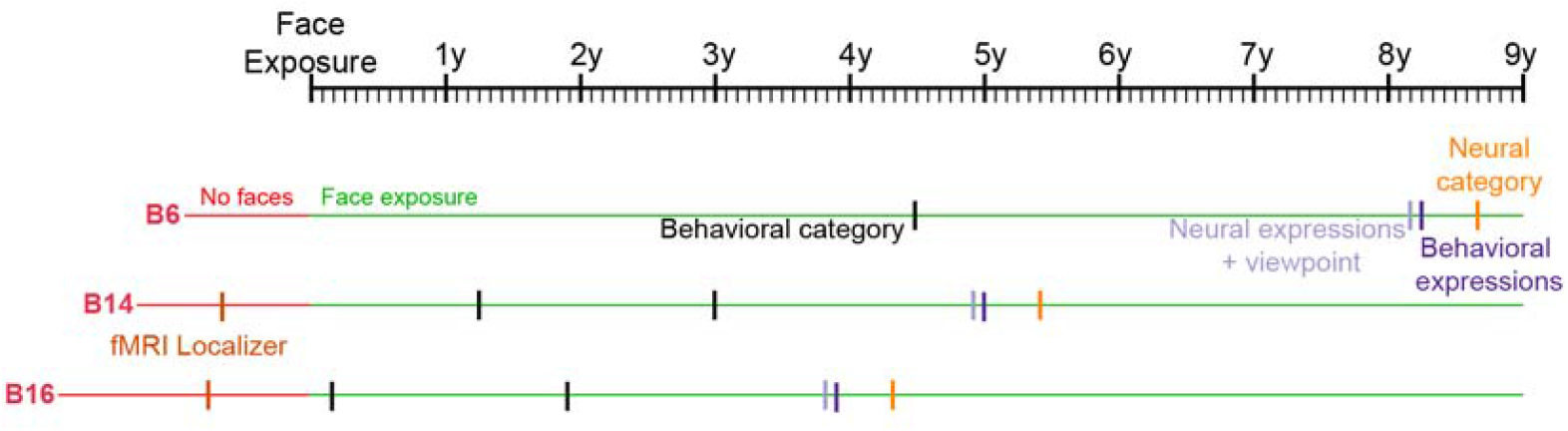
Experimental timeline for the three face-deprived monkeys (B6, B14 and B16). The experimental timeline aligned to face exposure. The results for the fMRI Localizer are shown in Supplementary Figure 2, for Behavioral category experiment in Fig. 1, for Behavioral expressions in Fig. 2, for Neural category in Fig. 3, and Neural expressions + viewpoint in Figs. 4 and 5.

**Supplementary Figure 2.**
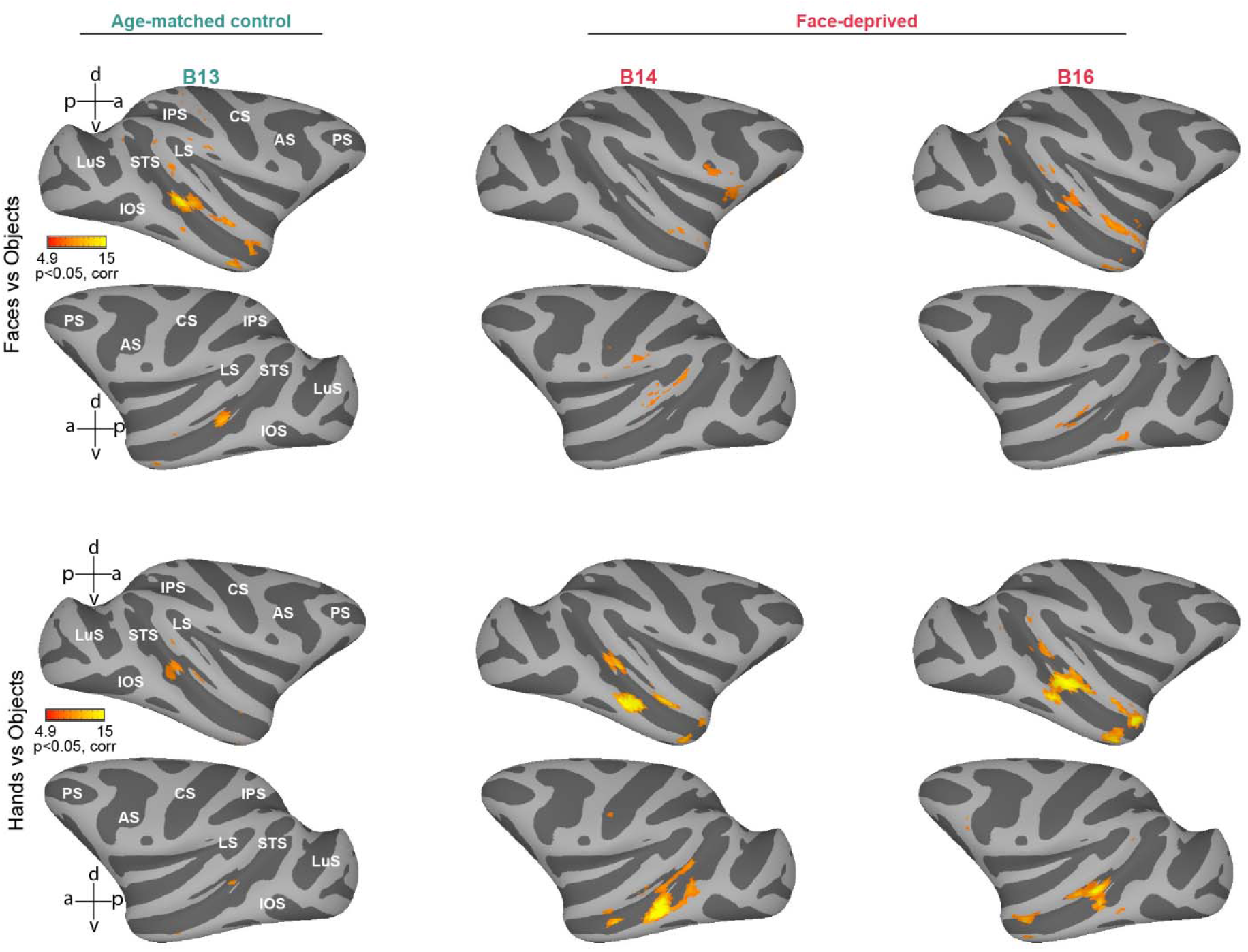
Categorical selectivity in face-deprived monkeys during the deprivation period. Heatmaps depicting fMRI activations on the inflated cortical surface of the standard NMT template for the contrasts Face > Objects (top two rows) and Hand > objects (bottom two rows) in two face-deprived monkeys and one age-matched control. Scans were acquired at 222 days old for B13 (control) and 214 and 382 days old for B14 and B16, respectively (face-deprived; before the end of the deprivation period). Orange shading indicates voxels with significant t-values (t>4.9; p<0.05, corrected). PS: Principal sulcus; AS: Arcuate sulcus; CS: Central sulcus; LS: Lateral sulcus; STS: Superior temporal sulcus; LuS: Lunate sulcus; IOS: Inferior occipital sulcus.

**Supplementary Figure 3.**
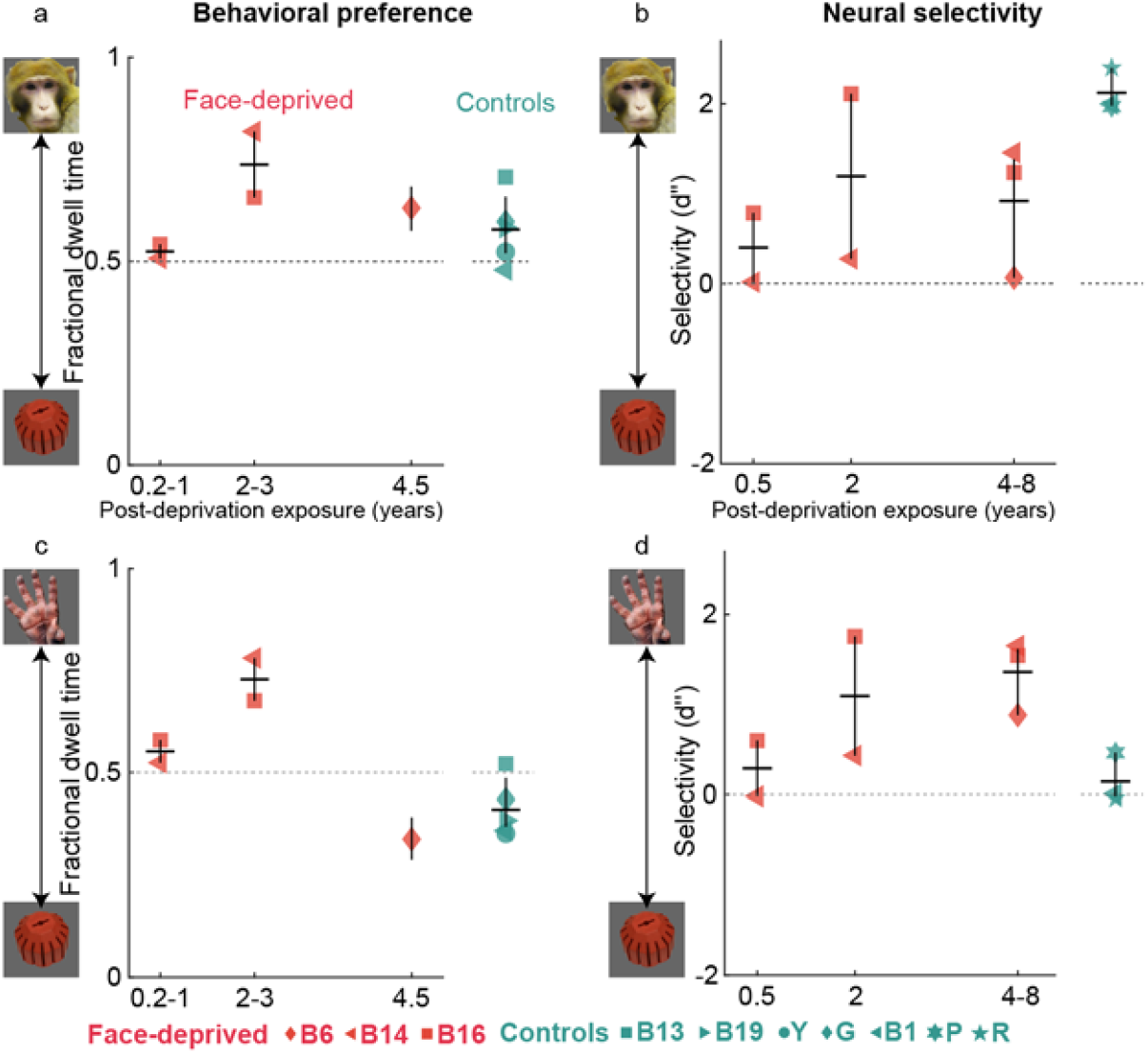
Categorical viewing preference and neural selectivity over time. **a**. Viewing data shown as fractional dwell time for faces vs objects for face-deprived (red) and control (teal) monkeys. Data for face-deprived monkeys is arranged chronologically by time after post-deprivation face exposure (0.2-1yr, 2-3yr, 4.5yr). Individual monkeys are indicated by different symbols. Vertical black line indicates mean and horizontal black line 95% CI across monkeys. For B6 (red diamond), the horizontal black line indicates 95% CI across trials. Grey dotted line indicates chance. **b**. Neural data shown as selectivity (d’) for faces vs objects for face-deprived (red) and control (teal) monkeys. Data for face-deprived monkeys are arranged chronologically by time after post-deprivation face exposure (0.5yr, 2yr, 4-8yr). Individual monkeys are indicated by different symbols. Vertical black line indicates mean and horizontal black line 95% CI across monkeys. **c, d**. Behavioral preference (**c**) and neural selectivity (**d**) for hands compared to objects. Same conventions as a, b.

**Supplementary Figure 4.**
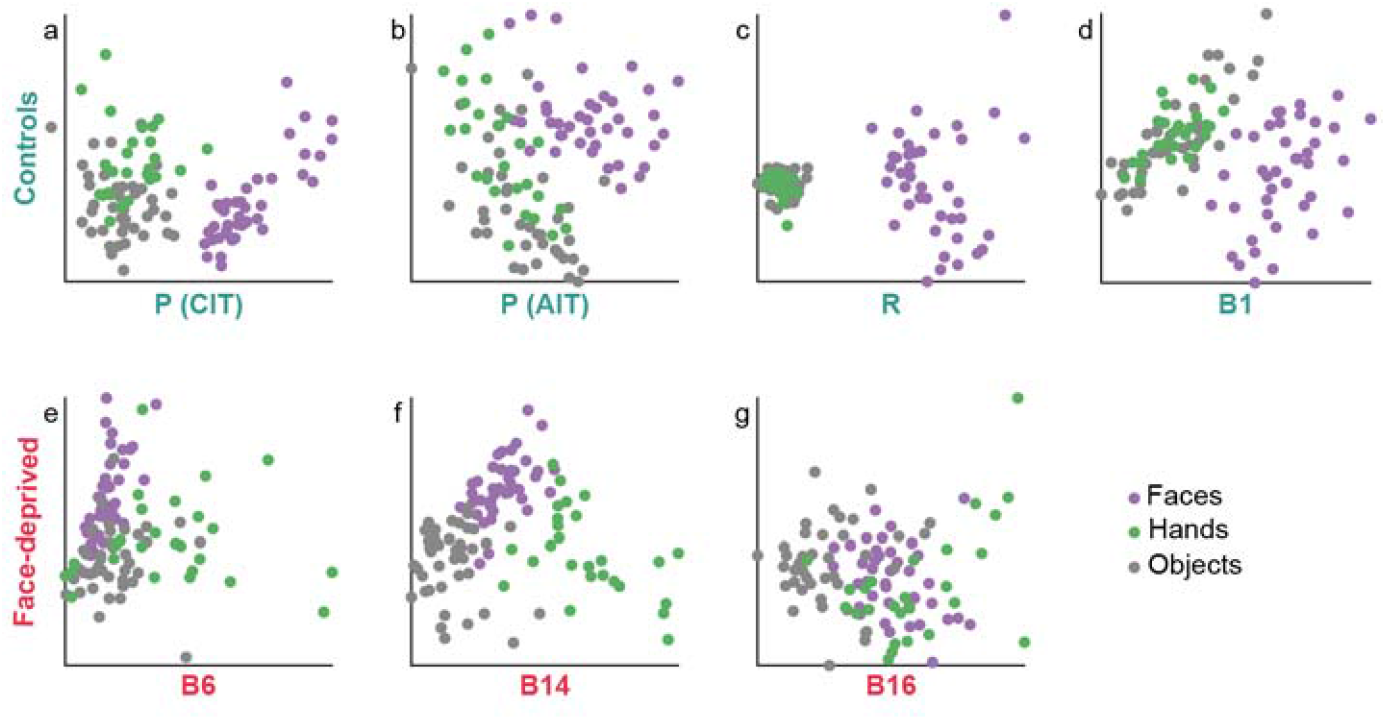
Representational structure of faces, hands, and objects across control and face-deprived monkeys. Multidimensional scaling (MDS) plots illustrating the representational geometry of trial-averaged population neural responses to faces (purple), hands (green), and objects (gray) for individual monkeys. Each point represents the population response to a single image, with distances in the plot reflecting the similarity of neural response patterns between images (see Methods). Thus, points that are closer together correspond to images that evoke more similar population responses. Top row shows control monkeys (P [CIT], P [AIT], R, B1), and bottom row shows face-deprived monkeys (B6, B14, B16).

## Acknowledgements

We thank Peter Schade and Dr. Michael Arcaro for assistance with the fMRI scanning. We also thank Dr. Kalanit Grill-Spector and her lab members for fruitful dicussions and feedback. This work was supported by the Harvard Lefler Fellowship and the Hearst Fellowship (to S.S.), NIH grants NS123778, P30 EY012196 and R01 EY025670 (to M.S.L.).

## Author contributions

Conceptualization: S.S. and M.S.L. Investigation: S.S. and M.S.L. Formal analysis: S.S. Methodology: S.S. Visualization: S.S. Supervision: M.S.L. Writing—original draft: S.S. Writing—review and editing: S.S. and M.S.L.

